# Open Science Saves Lives: Lessons from the COVID-19 Pandemic

**DOI:** 10.1101/2020.08.13.249847

**Authors:** Lonni Besançon, Nathan Peiffer-Smadja, Corentin Segalas, Haiting Jiang, Paola Masuzzo, Cooper Smout, Eric Billy, Maxime Deforet, Clémence Leyrat

## Abstract

In the last decade Open Science principles have been successfully advocated for and are being slowly adopted in different research communities. In response to the COVID-19 pandemic many publishers and researchers have sped up their adoption of Open Science practices, sometimes embracing them fully and sometimes partially or in a sub-optimal manner. In this article, we express concerns about the violation of some of the Open Science principles and its potential impact on the quality of research output. We provide evidence of the misuses of these principles at different stages of the scientific process. We call for a wider adoption of Open Science practices in the hope that this work will encourage a broader endorsement of Open Science principles and serve as a reminder that science should always be a rigorous process, reliable and transparent, especially in the context of a pandemic where research findings are being translated into practice even more rapidly. We provide all data and scripts at https://osf.io/renxy/.

## Introduction

The COVID-19 outbreak represents an urgent threat to global health. On October 15, 2020, the number of COVID-19 cases had exceeded 38 million and the death toll had exceeded 1,000,000 worldwide. Many important issues remain unresolved, including some crucial questions around both the diagnosis of patients with COVID-19 and optimal therapeutic strategies. Rapid scientific progress on these issues is needed to improve patient management, reduce mortality, and prevent new infections. The scientific community has responded accordingly, with the publication of over 80,000 preprints and peer-reviewed articles on COVID-19 or SARS-CoV-2 since announcement of the emergence of a new virus on 31^*st*^ December 2019 [1]. Many of these publications have contributed to the development of a body of knowledge that has since informed practice but a considerable number of these studies suffer methodological weaknesses, limiting the interpretability of their findings [2] or leading to false claims with a potentially dramatic impact on public health. While some of these studies have already been retracted [3, 4], others still contribute to the body of evidence and might be used by researchers and policy makers. In addition to the direct threat these publications pose to public health, these low-quality studies also exacerbate the waste of scientific resources [2] that is well-known to plague the scientific system [5]. Furthermore, many news outlets have recently amplified public exposure to low-quality research, sowing confusion among the public. In this paper we argue that many of the sub-optimal and non-transparent scientific practices witnessed during the pandemic, in conjunction with poor coordination across the global research community, have contributed to a dysfunctional scientific process for COVID-19 research. We support this view by providing results from an analysis of COVID-19 publishing data in recent months, including an analysis of reviewing times, conflicts of interests and misuse of non peer-reviewed material. We further argue that the widespread implementation of Open Science principles – known to increase the rigour, reliability and reproducibility of scientific results [6, 7, 8, 9, 10] – could help optimize research efficiency moving forward, and thus improve health outcomes and economic costs related to COVID-19.

Broadly speaking, Open Science aims to optimize scientific conduct and communication by exposing the scientific process, and results thereof, to the scientific community and broader public. This idea is implemented concretely through a number of core Open Science practices [8, 11, 12]: Open Access, Open Source, Open Data and Open Peer-Review. The best-known of those, Open Access, consists of making all scholarly communications freely available with full re-use rights. Open Access also encompasses early dissemination of manuscripts in the form of preprints (articles not yet published in scientific journals). Even though preprints are not yet peer-reviewed and thus could contain mistakes which may have been identified through an independent review process, they contribute to a more transparent and open scholarly publication system, accelerating reviewing and communication within the scientific community [13]. Open Source and Open Data aim at ensuring that materials such as questionnaires, forms, procedures, collected data, metadata, and source code are shared to foster replication studies, increase data re-use, and facilitate the peer-reviewing process [14][15]. Indeed, reviewers have the material at hand to verify the findings or detect any issues that could not be otherwise identified from the manuscript itself and to provide comprehensive peer-review reports. Then, following the Open Peer-Review principle, these peer-review reports should be publicly and transparently shared, along with the authors’ response. The scientific discussions between authors and reviewers are inherent to the process of creation of knowledge [16]. In addition, Open Peer-Review helps maintain high reviewing quality [17, 18, 19] and reduces the risk of concealed conflicts of interest. Therefore, the adoption of Open Science principles in the last decade has been particularly helpful in increasing the rigour, reliability and reproducibility of scientific results across research fields [6, 8, 9, 10].

There is evidence suggesting that the COVID-19 pandemic has served as a catalyst in the adoption of certain Open Science principles. For instance, major publishers such as Elsevier [20] and Springer Nature [21] have made newly written COVID-19 related articles freely accessible to all (Open Access). Furthermore, authors have shared their preprints more systematically than in previous pandemics [22] and reviews have been posted on external platforms (e.g., Pubpeer [23]). Specific initiatives, such as OpenSAFELY [24], have emerged to make data available to researchers while complying with the legislation regulating the use of medical data. Nevertheless, there have been many instances where these principles were ignored. One notorious example is the lack of transparency and sharing of the data provided by Surgisphere, which led to the retraction of the publication in The Lancet [25]. In other instances, some of the Open Science principles were adopted but misused. For example, news agencies have reported unreliable results based on unreviewed preprints [2] and some open reviews took place on separate platforms (for example Pubpeer), and were thus not directly available to readers.

While we recognize that the faster embracing of Open Science during the pandemic is a step towards more accessible and transparent research, we also express concerns about the adoption of these practices for early and non-validated findings. Furthermore, embracing only some of these principles, while excluding others can have serious unintended consequences that may be as detrimental as not adopting open practices in some instances. The aim of the present paper is twofold. First, we identify the issues the scientific community has faced with regard to the publication process since the beginning of the pandemic. To do so we analyzed data collected on preprints and published COVID-19 research articles, as well as on retracted COVID-19 publications, in order to quantify issues related to reviewing time, conflicts of interest, and inappropriate coverage in the media. In light of this analysis, we then discuss how a wider adoption of Open Science principles could have potentially minimized these issues and mitigated their impact on the scientific community and broader public.

The structure of this article follows the stages of the publication process shown in Figure 1. We first discuss issues arising at the data collection and interpretation stage (before the dissemination of the results). Then, we review the dysfunctions observed during the publication process (between the submission and the publication of research articles), before investigating the misuses of research outputs during science communication (after publication). We provide recommendations based on Open Science principles for each stage of the publication process, which we hope will contribute to better research practices in the future.

**Figure 1:**
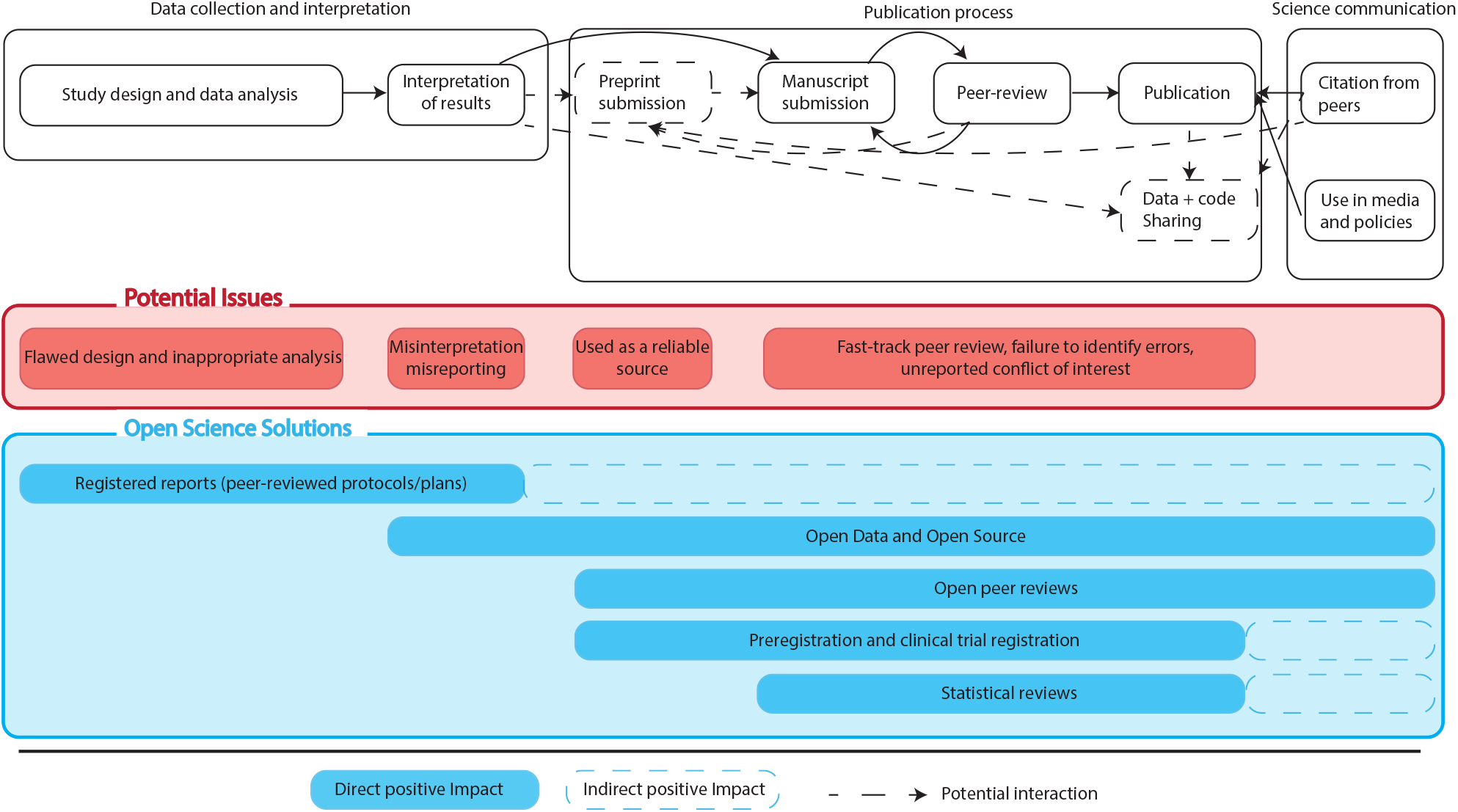
Outline of the publication process with its potential issues and our proposed solutions.

## 1 Stage 1: Data collection and interpretation

While previously deplored [5], waste of scientific effort has been particularly prominent during the COVID-19 pandemic [2] and has been more visible than ever before. In this section, we show that this waste has its roots in the early stages of the research process – at the data collection and interpretation stage – and discuss how study preregistration, registered reports, adherence to reporting guidelines and Open Source principles could help to minimize waste in research.

### 1.1 Identified flaws

#### 1.1.1 Methodological and statistical issues

Conducting research during a pandemic is known to pose particular challenges [26] but scientists have raised concerns about methodological flaws in the design and analysis of various COVID-19 pharmacological studies [2, 27, 28, 29].

To better understand whether inappropriate study designs or statistical analyses contributed to the reasons behind the retraction of articles, we looked at the 29 COVID-19-related papers that had been retracted (or are subject to expressions of concerns) since January 2020 [3]. The list of articles (both preprints and peer-reviewed) and the results of our analyses are available on the repository of this project: https://osf.io/renxy/. Of the 29 identified publications, 8 (27.6%) were retracted (or had an expression of concern from the editorial board) based on their data analysis or study design. More specifically, among these 8 publications, 2 (25.0%) papers were retracted, at the authors’ request, in order to conduct further data analyses and 6 (75.0%) were retracted because the methodology or the data analysis was wrong.

#### 1.1.2 Duplication of research

Another concern is the increased risk of research waste due to duplication. Many studies that aimed to assess the efficacy of hydroxychloroquine were conducted in parallel: 218 registered trials were ongoing or already completed as of 26^*th*^ April 2020 [30]. Many comparative effectiveness studies – randomised or not – were conducted without preregistration (e.g., Geleris et al. [31]), however, meaning that the broader research community only became aware of these studies at the time of the release of the results. This illustrates the general lack of cooperation between research teams, putting more patients at risk by exposing them to potentially harmful treatments in multiple underpowered studies, and also leading to a waste of research time and financial and human resources [2]. Given the additional workload for healthcare workers and clinical researchers these trials require, it may have contributed to the disruptions in the conduct of clinical trials during the pandemic [32]. Other studies have been pre-registered but conducted and reported with major deviations from the preregistration record without justification (e.g., [33]).

#### 1.1.3 Ethical concerns

Ethical concerns have also arisen during the pandemic. While the research community needs to find ways to provide timely solutions to the COVID-19 crisis, it should not be at the detriment of good research and clinical practice. Among possible ethical risks, Xafis et al. [34] identified over-recruitment in trials, the conduct of human vaccine studies before the completion of animal studies, and the neglect of adverse effects in drugs studies. An example of the last is the little consideration given to the known cardiotoxicity of the combination of hydroxychloroquine and azithromycin early on in the pandemic. Issues surrounding patients’ participation in clinical studies have also been observed: in her analysis of COVID-19 papers unsuitable for publication, Bramstedt identified issues surrounding informed consent as the second most common source of concerns [35]. In addition to the ethical problems this poses, it could also weaken the trust that patients and the broader community afford researchers, with detrimental consequences for public health in the long term.

### 1.2 Open Science solutions

Here, we argue that the adoption of certain Open Science principles could have helped to detect or avoid the issues in data collection and interpretation described above (1.1). Two methods seem to be particularly relevant:

#### 1.2.1 Study preregistration

First, study preregistration on dedicated platforms (e.g., ClinicalTrials.gov, [OSF](https://osf.io/), or [AsPredicted](https://aspredicted.org/)), with a thorough description of the study design, ethical approval, methods for data collection and data analysis, can help prevent some of the issues identified above (1.1). Indeed, study preregistration may reduce the amount of unnecessary duplication of research as researchers will be able to check whether specific studies are ongoing and design theirs to address complementary questions. Finally, study preregistrations can be used by Institutional Review Board (IRBs) for ethics approval and to fulfil the ethical obligation to transparently inform both the public about ongoing trials as well as the research community [36].

Another goal of preregistration is for readers and reviewers to make sure that a published study has been conducted and analysed as planned, thus limiting the risks of changes to the design, methods or outcomes in response to the data obtained other than the flexibility allowed by the protocol (in case of interim analyses of adaptive designs). Researchers should register studies prior to data collection. On the platform Clinical-Trials.gov, retrospective registrations or updates to the study protocol are flagged. Depending on the level of methodological details in the record, study preregistration may help in limiting questionable research practices such as HARKing [37], p-hacking and p-fishing [7] and eventually lead to better subsequent reporting [38].

As COVID-19 was a new disease, there was no standardized diagnostic criteria or clinical outcomes. This led to a multiplication of different outcomes studies in the articles participating in the difficulty to replicate and compare results. Study preregistration could help researchers adopt the same criteria and outcome measurements and promote the use of validated international standardized criteria for variable and outcome measurements [39].

However, such preregistrations have two major limitations. First, they do not fully prevent duplication. While replication (defined as a deliberate effort to reproduce a study to validate the findings) is an important step of the research process, duplication (an inadvertent repetition of the research) contributes to research waste. This waste has been noted among COVID-19 research [2], with a strikingly high amount of duplication despite study preregistration. Second, whereas preregistrations allow the detection of questionable research practices, they do not help prevent methodological issues before data collection since the preregistration is not itself peer-reviewed and the statistical analysis section of these records is often very brief. Therefore, study preregistrations are necessary, since they encourage researcher to outline the study design and analysis strategy, but not sufficient to avoid the excessive waste of scientific resources.

#### 1.2.2 Registered report

Peer-reviewed study protocols, also called registered reports [40, 41], can also have a major impact on the reduction of wasted resources. Registered reports can be adopted in any research area using experimentation. They essentially consists in articles with a two-stage peer-review, and provide details about the research question, hypotheses, methodology, statistical analyses and reporting strategy. Since protocols are peer-reviewed before the enrollment of participants and data-collection, potential omissions or mistakes in the proposed methodology can be corrected before any substantial resources are used, thereby limiting scientific waste [5, 42]. In these reports, researchers are also encouraged to provide details about the resources used, using for instance Research Resource Identifiers (RRIDs) [43] when applicable, and specify the reporting guidelines that will be used (e.g., CONSORT [44], STROBE [45]). Registered reports can therefore contribute to higher quality research, with a reduced risk of bias and increased generalizability. These improvements in quality and robustness of scientific evidence [7] ultimately facilitate its communication and use. One disadvantage of registered reports is that their reviewing takes time, while preregistrations are immediately available. However, some platforms for the submission of registered reports put in place measures to guarantee a timely review of COVID-19 protocols: stage 1 review of registered reports at Royal Society Open Science are performed within 7 days [46]. Furthermore, since they reduce the risk of publication bias, they also reduce the number of submissions needed to publish one’s results and should ultimately save both resources and time (e.g., [47]). Both pre-registration and registered reports contribute to a better visibility of ongoing research, and should be used at institution level to coordinate research projects at an international level in a more efficient way, in order to optimize resources.

#### 1.2.3 Beyond registration: Open Methodology and reforming the publication system

Preregistrations and registered reports are necessary but not sufficient to conduct reliable transparent research. The principle of Open Methodology [12] therefore goes further. It consists in transparently sharing all the necessary details to allow replication of the research. In other words, Open Methodology relies on the authors to not withhold any details of their research project so that any outsider could exactly replicate it. While one might assume that preregistrations and registered reports are sufficiently detailed to allow replications, numerous past studies have shown that replicating research work is excruciating or impossible. Journals should therefore support fully Open Methodology better by enforcing that all submitted articles must be fully reproducible before the manuscript is published. Beyong Open Methodology and registrations, researchers have suggested to reform scientific communication so that they would be less story-telling oriented and more focused on the methodology (see Octopus [48]). Every step of research methodology (research-question formulation, hypothesis-making, data-collection plan, data analysis, interpretation…) is a smaller paper that builds onto the previous one and all of them are open for comments and reviews in order to make the insights more robust and foster collaborations. Adopting this would, however, require a complete change of the scientific publishing system and further proof is needed to show its benefits.

### 1.3 Section conclusion

In clinical research, the adoption of study preregistration for clinical trials (called clinical trial registrations) and observational studies, as well as registered reports, have contributed to better transparency and reliability of the findings. For the researchers themselves, the availability of the report facilitates complete and transparent reporting in the final publication. Finally, study preregistration and registered reports enable the conduct of more exhaustive meta-research and systematic reviews, by reducing the risk of publication bias. We therefore argue that both study preregistration on dedicated platforms, already mandatory for some types of clinical trials, and registered reports should be used more systematically across research fields.

## 2 Stage 2: Publication process

Publishing scientific evidence consists of several steps summarized in the second box in Figure 1. In response to the COVID-19 global pandemic, an enormous number of research publications have been produced, both in the form of preprints and peer-reviewed articles. Unfortunately shortcuts have been taken in the publication process of some of these papers, jeopardizing the integrity of the editorial process and putting the rigour of scientific publications at risk. In this section we discuss three issues in the peer-reviewed publication process which have arisen during the COVID-19 pandemic: fast-track publication, conflicts of interest and lack of data sharing. We highlight the increasing retraction rate of COVID-19 preprints and peer-reviewed articles, and propose practical solutions to minimize these issues in the future.

### 2.1 Identified flaws

#### 2.1.1 Expedite reviewing and conflicts of interests

The publication pipeline has been directly affected by COVID-19, in particular with respect to reviewing times of COVID-19 papers [49]. Although fast-tracking particular articles is not uncommon in the scientific publishing system [49], a number of journals have recently implemented specific policies to fast-track COVID-19-related research (e.g., PLOS, some Wiley journals [50], some Elsevier Journals [51], some SAGE journals [52], and PeerJ journals [53]). In addition a new overlay journal for fast and independent reviews of COVID-19 preprints has recently been launched [54]. While faster peer-reviewing does not necessarily equate with poorer review quality and faster peer-reviewed time are encouraged at a time of crisis, it remains unclear how thorough the peer-reviewing is and how potential conflicts of interest are handled. Palayew et al. [49] recently highlighted that COVID-19-related manuscripts between 1 January 2020 and 23 April 2020 had a median reviewing time of 6 days and that more than a thousand manuscripts were reviewed in less than 7 days.The authors also identified manuscripts for which it was unclear whether they had been reviewed at all. More recently, Homolak et al. compared median submission-to-publication times of COVID-19 publications with those of non-COVID papers during the same period and before the pandemic [55]. These times were reduced by a factor 10 to 15 for COVID-19 papers, consistently across research fields. It should be noted that the FASEB Journal, whilst acknowledging the degree of risks in doing so [50], allows editors to directly accept COVID-19-related submissions for Review, Perspectives, and Hypotheses, without peer-review, as per the journal’s fast-track policy [56]. In light of this, we have sought further information on the fast-tracking of peer-reviews with up-to-date information.

We searched for “COVID-19”(and related terms, see the full list in Appendix) on PubMed Central and found 12,682 published articles as of 1^*st*^ July 2020. Of these we could extract the reviewing time for 8,455 (66.7%) articles, as the difference between the date of submission and the date of acceptance. Of these 8,455 publications, 699 (8.3%) from 341 different peer-reviewed journals were reviewed and accepted for publication either on the day of submission (*n* = 311) or the day after (*n* = 388). We manually inspected these manuscripts to identify potential conflicts of interest. We focused only on editorial conflicts of interest, i.e., manuscripts for which at least one of the authors is an editor-in-chief, associate editor or a member of the editorial board of the journal in which the article was published. While publishing articles with editorial board members among the co-authors is not problematic in itself, and is often encouraged by publishers provided that a specific process is put in place [57], we were concerned about the lack of transparency in the peer-review process of these articles. We did not assess institutional or financial conflicts of interest. Among the 699 articles accepted within a day, an editorial conflict of interest was observed in 297 (42.5%) articles. In order to assess whether the frequency of these conflicts decreases over time, we also inspected the manuscript accepted in 16 days and 20 days, corresponding respectively to the median and mean time between submission and acceptance in our sample. To further investigate the distribution of these conflicts, we divided the articles into two groups: research articles, defined as publication with original findings and other types of publications, including editorials, letters and reviews (without a pre-defined methodology. The proportion and type of conflicts of interest per type of article and time between submission and acceptance are presented in Figure 2.

**Figure 2:**
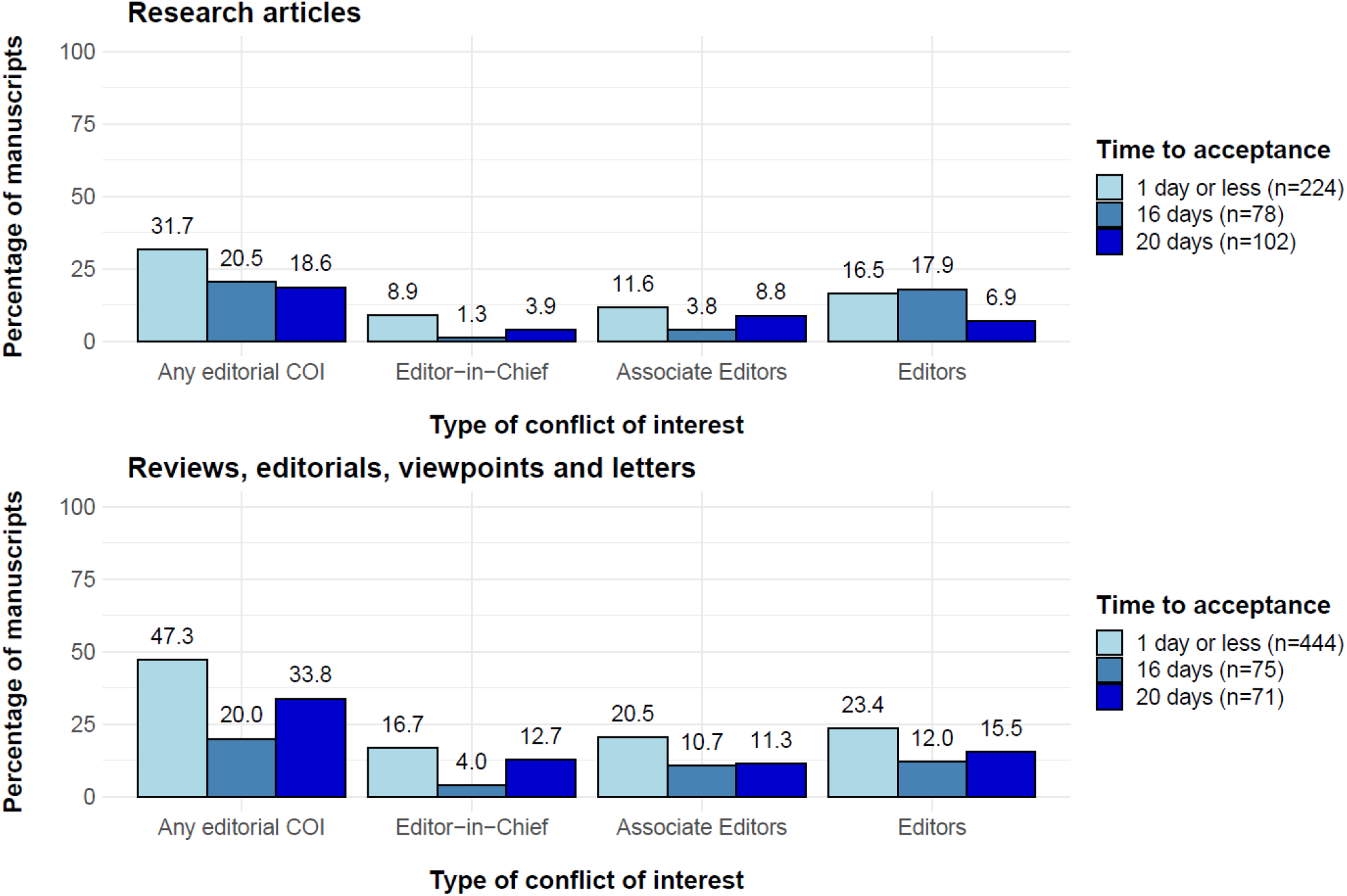
Distribution of conflicts of interest according to the type of article for COVID-19 research articles with a submission-to-acceptance time of a day or less, 16 days and 20 days. COI: conflict of interest. Note: for fairness of comparison, we restricted our analysis to articles submitted before 11^*th*^ July 2020, since it was the last submission date at which an acceptance time of 20 days could be observed.

As expected, conflicts of interest were most common for editorials letters and reviews, but were also surprisingly frequent for research articles (*n* = 71, 31.7% among articles accepted in a day or less). The prevalence of these conflicts was substantially heterogeneous across journals: the estimated intraclass correlation coefficient for the proportion of publications with any conflict of interest was 0.37, (95% confidence interval: [0.29; 0.45]), which means that 37% of the variability observed in the occurrence of conflict of interest can be explained by the journal in which the articles were published. The frequency of editorial COIs decreases with the increase in time to acceptance for both types of papers, although there are still common at 20 days (*n* = 19, 18.6% for research articles, *n* = 24, 33.8% for other types of manuscript). Unfortunately, it was not possible to perform the analysis for papers with longer time to acceptance, as it would have restricted the analysis to papers submitted in January and February 2020, reducing the sample size and the generalizability of the results. These findings raise concerns about the fairness and transparency of the peer-review process with such short acceptance times. For example, an opinion paper, reviewing the literature on Hydroxychloroquine and Azithromycin as an early treatment for COVID-19, written by a member of the editorial board of the journal, has been published in the American Journal of Epidemiology within 7 days of submission [58]. This article was followed 3 months later by an expression of concerns from other members of the editorial board, who identified major flaws in the review [59].

While the need for faster scientific dissemination during a pandemic is understandable, the possibility to publish without a rigorous and critical peer-review process is, in some circumstances, detrimental to the scientific community and the public at large. This is the case when these findings are used to inform medical practice or public health policies. For example, following concerns about the scientific validity of a study investigating the effectiveness of hydroxychloroquine, accepted for publication in less than a day after submission [33], post-publication reviews were commissioned by a learned society (International Society of Antimicrobial Chemotherapy). These reviews, published 4 months after the initial publication, [60, 61], pointed out major methodological and ethical flaws. Despite this, the paper was not retracted, on the grounds that it gives room for scientific debate [62]. This is problematic for several reasons. First, such papers with dangerous conclusions are still available to researchers, with no mention within the article of the existence of the post-publication reviews as a warning. Secondly, this type of study is likely to be included in any subsequent systematic review done on the topic, and, even though it may be flagged as a study with a high risk of bias, it might influence the results of the meta-analysis. Finally, the choice of the editorial board not to withdraw this publication, but instead encourage the submission of letters or comments, will help increase the impact of the journal, despite the poor quality of the original publication and associated review process.

#### 2.1.2 Distrust of published results

In 2016,, identified a phenomenon that had later been called the ‘reproducibility crisis’ [63]. In this survey, more than half of the respondents reported that they had experienced trouble reproducing published results (including their own) at least once, raising doubts as to the quality of these published results. This issue has two main causes: the lack of data sharing and the lack of code sharing.

At the heart of biomedical research lies the need for high-quality data. From these data, using a suitable methodology and statistical tools, researchers can answer relevant clinical questions. While privacy issues raised by collecting medical data should be addressed, there is a need for data sharing between researchers to reproduce the results, enhance collaboration and obtain more timely results. Achieving both of these goals, however, remains a challenging task for the scientific community, especially during an emergency.

To date, four peer-reviewed articles related to COVID-19 have been retracted shortly after publication due to concerns about potential data fabrication or falsification. Two retracted articles, in particular, attracted a great deal of attention: one in the New England Journal of Medicine [64] and one in The Lancet [25]. These articles reported the findings of an international study, based on data owned by the company Surgisphere. The data were not publicly shared at the time of submission or after publication but, even more disturbing, the data were not shared with all the co-authors. The initial publication of the study in The Lancet [25] indicated that hydroxychloroquine increased mortality for COVID-19 patients. These serious safety concerns led the WHO and INSERM to interrupt the inclusions in the hydroxychloroquine arm of the Solidarity and DisCoVeRy trials ([65, 66]) while the clinical data of patients treated with hydroxychloroquine were reviewed. Our review of the retracted COVID-19 papers highlights that two additional papers using Surgisphere data have also been retracted. Surgisphere’s refusal to share data with the scientific community, including authors involved in the study, and even to a third trusted party, eventually led to the retraction of both articles. The most striking consequence of this affair, however, is that it may have made scientists, editors, readers, organizations and reviewers waste precious research time, when a rapid response is needed.

### 2.2 Open Science solutions

#### 2.2.1 Open Review

As previously stated, we understand and acknowledge the pressure and the need to accelerate reviewing of submitted manuscripts, but journals and editors should carefully consider the trade-off between reviewing quality and reviewing time. Our findings relating to the fast-tracking of peer-reviews for COVID-19-related articles in Section 2.1.1 can, however, be perceived as particularly worrying. In a time of a pandemic, medical management of patients and public health policies rely heavily on scientific findings. Fast-tracking of peer-review should therefore only be done when scientific rigour can be maintained as its loss might lead to disastrous consequences for patients and for public health as a whole. A greater transparency in the peer-review process is thus urgently needed. Sharing reviewers’ reports along with the authors’ response more systematically could contribute to this transparency. These scientific discussions are extremely valuable as they may help balancing the views expressed in the published article. In addition, reviews are usually found to be of higher quality when they are made publicly available [17, 18, 19]. The availability of the reviews may also help the scientific community and the different stakeholders to verify that the peer-review of a manuscript has been thorough and so help to increase public trust in scientific research. Several implementation of Open Reviews exists and the name is loosely used to define all of them [17]. In the present case, we argue that making reviewers’ reports available, should become the norm to help mitigate the issues previously highlighted in subsubsection 2.1.1, but as correctly argued in the literature (e.g., [17]) enforcing signed open peer-reviews might be problematic, although it has been adopted by some journals.

Furthermore, the peer review process should evaluate all aspects of the publications, in particular the methodology, in order to identify inappropriate study designs or incorrect use of statistical methods (see subsection 1.1). In order to do so some journals have decided to resort to specialist statistical reviewers [67]. The British Medical Journal’s editorial board, for instance, includes a Chief Statistical Advisor and Statistics Editors [68]. The reviewing process is conducted by domain experts and statisticians in order to make sure that the claims or findings hold with respect to the conducted statistical analysis but also to ensure that the statistical analysis conducted by the authors is appropriate and correct. Statistical reviews help in the detection of mistakes in the data analysis and in detecting exaggerated claims in a manuscript before publication. A recent review of retracted research articles showed that among the 429 papers that were retracted from journals in which statistics are assessed, 81 (18.9%) had a specific statistical review, whereas the assessment of statistics was part of the reviewer or editor’s task in the remaining 348 (82.1%) [69]. Although the retraction rate was lower among articles involving a specialist statistical reviewer, the low rate or retraction prevents us from drawing any strong conclusions (5 per 10,000 vs 7 per 10,000).

To better anticipate, detect, and act upon the potential issues raised in subsubsection 2.1.1 about fast-tracking or conflicts of interest, but also to detect studies that do not reach the expected standards for research, we believe that complete transparency in and around the peer-reviewing process should be adopted. In light of the lessons learned during the current pandemic we therefore recommend the following to be adopted by journals and editors:

1. Authors should highlight, in their manuscript and its metadata, any affiliations with the editorial board of the journal to which they submit their paper. An example of such disclosure can be seen in Pardo et al. [70].
2. Journals should explicitly state how the peer review was conducted: they should state how many referees were recruited and the duration of the complete review process. This should include how long it took to find referees, how long each referee took to complete the review (time between submission and acceptance of the paper) and the number of iterations between the reviewers and the authors.
3. In addition, journals should make the referees’ reports of accepted articles transparently available alongside the manuscript allowing authors, reviewers and the scientific community to benefit from the constructive comments in these Open Reviews. Journals should probably also consider the various Open Review processes [17] and let referees themselves decide whether or not they want to sign their reviews.
4. When publications report quantitative findings, a systematic review of statistical methodology should be included.

Although the peer-review process implemented by journals contributes greatly to the quality of manuscripts, each submission is typically reviewed by a limited number of reviewers whose skills might be too specific to evaluate the different components of a manuscript. Crowdsourced peer-reviews (a.k.a portable peer reviews) reviews that are not ‘on request’ but carried out spontaneously, whether on preprints or post-publication. They are complementary to the journal’s peer-reviews. They are open because they are available to all and, very often, signed. The use of these peer-reviews through dedicated platforms such as Pubpeer [23] or Zenodo [71], or even ASAPbio[72], Review Commons[73] and the COVID-19-pandemic-born OutbreakScience PREreview [74] that all allow solicitating reviews on preprints, has gained popularity during the pandemic. For instance, statisticians have published a detailed and comprehensive review of the preliminary report of the RECOVERY trial: the largest comparative effectiveness study on COVID-19 treatments to date [75]. Further examples are presented here [76]. It is notable that the reasons for the retraction of 5 COVID-19 papers were echoed on the Pubpeer link to those publications. Therefore, open reviews can contribute to an early detection of flaws in research articles. However, providing a thorough feedback on publications is time-consuming and reviewing activities are usually not highly valued by institutions, nor strongly encouraged. Therefore, these changes in peer-review practices cannot be implemented without meaningful support and endorsement from research institutions. Many institutions have created repositories of accepted manuscripts to promote Open Research. These initiatives should go a step further by encouraging researchers to archive the reviewers’ reports as well.

We postulate that adopting Open Reviews and complete transparency in the reviewing process, in addition to all the already highlighted benefits identified in the literature [17, 77, 18, 19], would have helped in detecting potential mistakes in manuscripts or frauds and saved precious research time during the pandemic. Furthermore, it would make analyses of peer-review processes and their quality easier to conduct [78] and would accelerate the training of Early Career Researchers [78, 79] whose help might be critical during a public health crisis.

#### 2.2.2 Open Data and Open Source

To aid a more thorough peer-review, enable data re-use, and facilitate reproducibility, researchers are now being encouraged to share their data on publicly accessible repositories [80, 81], along with the code used for the analysis. Even though sharing raw data may be challenging in medical research due to the need for compliance with data protection regulation, Open Data initiatives have been considered one of the main solutions to avoid a replication crisis [82, 15] and are often seen today as a critical part of the peer-review system [15].

Data sharing, including anonymized raw clinical data, is crucial during a pandemic and could accelerate the understanding of new diseases and the development of effective treatments [83]. Several researchers have thus, early-on, argued that journals and institutions alike should always ask authors of manuscripts to confirm that the raw data are available for inspection upon request (or to explain why they are not) [84, 85]. Considering that data fabrication and/or falsification has been observed in the scientific community and its frequency probably underestimated [86], some have suggested [15] that the policy of “sharing data upon request” is not enough. Therefore, recent years have seen an increase in policies, from journals and institutions alike, asking researchers to share their raw data by default, except when data could not be shared for privacy reasons [82, 15]. While data sharing is increasingly being adopted, it still does not appear to be a default practice [15]. Maintaining a high-quality database is time- and money-consuming but sharing it allows other researchers to conduct research with the data and optimize the costs. The MIMIC database, a freely accessible critical care database, has led to thousands of articles using and citing this database, and increased clinical and scientific knowledge in this field [87].

Another approach to the reproducibility crisis is making the source code used for the analysis openly available to the public and the scientific community [88]. Solutions have been proposed by the research community to promote and facilitate code sharing. An open code repository has been created [89] and many scientific journals are now asking researchers to publish the source code alongside their findings. The use of source code sharing platforms, such as GitHub or open source alternatives such as GitLab, is becoming common and has even been advised to improve open science behaviour [90, 91, 92]. The code should be published under a free license to encourage re-use and further developments, and when possible, open source software and programming languages should be preferred to maximize accessibility and reproducibility. The lack of code sharing is still, however, a major issue in the biomedical literature. In times of crisis, the need for open code is even more crucial when scientific evidence is at the root of major political decisions.

One may argue that the four retracted papers relying on Surgisphere data would not have been published if proper data sharing policies were in place for all journals. The policies currently in place, while already effective in reducing scientific fraud [15, 84, 85, 93, 94], are not sufficient to detect the issues that the Surgisphere papers have raised. Indeed, it is currently considered acceptable for authors to state that they cannot share their data (or code) for legal reasons (e.g., privacy [95]). We therefore urge that data-sharing policies should be adapted to the following:

1. Research material for data generation or data collection (e.g. research questionnaire, simulation algorithm, etc.) should be made available on dedicated platforms, and when appropriate published under a CC-BY Creative Commons license.
2. Data should be shared by default: authors should not be able to submit a manuscript if they do not provide access to raw data and analysis scripts or a valid reason why they think it is not feasible. This is already the process used by some journals: Wiley has clear data sharing policies, and data sharing is mandatory for submission to some of its journals (e.g., Ecology and Evolution) [96].
3. If authors are not able to share their raw data, journal editors should be able and should strive to demand that raw data be examined by a trusted third party (not the authors’ institution) to establish the existence of the raw data and validate the results of the analysis presented by the authors. Identification of the trusted third party can be left to the discretion of the ethics committee or the proprietary company but should not present any conflict of interest with the authors of the manuscript. The trusted third party should produce a public and signed report stating that the data are available and that the results presented in the manuscript can be confirmed.
4. To facilitate meta-analyses, abstracts of all manuscripts should contain links to preregistration numbers, data repositories and open source repositories. Additionally, publishers should consider directly including such information in paper’s XML to allow for even easier retrieval through text mining.

While we understand that changing data-sharing policies will take time, we hope that the scientific community, data-holding companies and governmental agencies protecting rights to privacy will learn the lessons from this pandemic and consider adapting their policies to the aforementioned points.

## 3 Stage 3: Science Communication

Once a manuscript is published, it is available for the rest of the scientific community to cite or conduct meta-studies on, but also to be communicated on by the media or to support policy makers. In this section, we review some of the issues that the pandemic has highlighted with respect to media coverage of scientific results following the publication of preprints and peer-reviewed articles, and we propose alternatives for a more responsible communication of scientific findings.

### 3.1 A surge of preprints and their misuse

Preprints are articles that authors make available to the community before they have been peer-reviewed, usually around the time of submission to a peer-reviewed journal. There are several benefits of preprints. They allow the communication of new findings to the research community in a more timely manner [13, 97], especially in the context of an emergency such as the current COVID-19 pandemic. Although these results should be critically assessed by the readers, having not yet received the benefit of peer-review, they might encourage the conduct of replication studies or further research building on these findings. Preprints also contribute to the reduction in wasted research by signaling ongoing research projects, avoiding duplication and potentially fostering collaborations. Preprints may also increase a researchers’ visibility and help in credit attribution and priority of DisCoVeRy [13, 98], reducing the risk of plagiarism. In addition, they are an opportunity for authors to get early feedback from a wider range of researchers and incorporate the suggestions received to enhance the quality of the publication [13, 98].

The COVID-19 pandemic has seen a surge in the numbers of preprints submitted by researchers [99]. While 807 preprints were deposited on medRxiv in the six-month period between 1^*st*^ July 2019 and 31^*st*^ December 2019, 6,771 preprints were submitted in the next six months (between 1^*st*^ January 2020 and 30^*st*^ June 2020), an increase of 739%. These figures are, respectively, 15,838 and 21,804 for bioRxiv (38% increase) and 87,942 and 112,197 for arXiv (28% increase). A recent study [100] gives more insights on the scale of this disruption. It has been found that 32% of the COVID-19 articles from the NIH OPA iSearch COVID-19 Portfolio [101] were preprints while, on the same time window in 2019, there were only 3% of preprints in the biomedical literature [100]. The use of preprints during outbreaks is certainly not new: a systematic review identified the publication of 174 and 75 preprints during the Ebola and Zika virus outbreaks, respectively [102]. Nevertheless, these figures are much smaller than the number of preprints submitted in the first 6 months of the COVID-19 pandemic. Consequently platforms offering preprint hosting have had to rapidly adapt to the rate of submission and adjust their screening procedures for each submission to avoid the dissemination of misleading or blatantly false claims [99].

Although the surge in preprints during the pandemic can be seen as an encouraging step towards Open Research, preprints – by their very nature – contain unreviewed findings. Although scientific findings from a single study should always be interpreted with caution, the interpretation of results from preprints invites even further caution. Unfortunately, some COVID-19 preprints have been misused, notably regarding communication with the general public. Indeed, various news outlets and also scientists have used these non peer-reviewed articles as scientific evidence, increasing the impact of potentially inaccurate findings. While external peer-review does not guarantee the validity of the results and comes with its own limitations, it contributes to the improvement of the quality of research outputs. However, the number of reviewers per manuscript is typically low (2.2 on average in a review done by Huisman et al. [103]), which may leave gaps in the expertise required to review all aspects of the articles. Thus, one of the benefits of preprints is to receive feedback from many other researchers with a broader range of expertise, which helps to identify and correct potential flaws in the methodology, analysis or reporting, thus enhancing the quality of the article prior to submission. As such, preprints may contain inaccuracies or unreliable findings and it must be noted that many preprints are never accepted for publication in peer-reviewed journals [104]. Looking at previous pandemic data in particular, a systematic review of preprints during the Ebola and Zika outbreaks [102] highlighted that only 48% of the Zika preprints and 60% of the Ebola preprints could be matched with peer-reviewed publications that later appeared. Although we cannot exclude the possibility that the authors never submitted their preprints to a peer-reviewed journal, a potential explanation for this phenomenon could be the presence of concerns expressed by the community on the preprints, highlighting poor methodology or other flaws that rendered the preprints unsuitable for publication.

In order to estimate how often preprints platforms were mentioned in the media, we queried the Factiva news database using the three major platforms hosting COVID-19-related preprints (arXiv, medRxiv and bioRxiv). On the 13^*th*^ of July 2020 the three platforms had been mentioned a total of 3,288 times in online news and in 313 blog posts since 1^*st*^ January 2020, the day after China alerted the WHO about a newly identified virus. Looking at English-only shares, we find 2,193 web news items reporting scientific findings from recent preprints and 121 articles addressing the surge of preprints during the pandemic and its challenges. These findings raise the concerns that the public may not have been correctly informed about the preliminary nature of these findings.

Next, to quantify the extent to which preprints themselves were shared in the media (news media and social media), we conducted a systematic search for preprints submitted to arXiv, medRxiv and bioRxiv between the 1^*st*^ January 2020 and the 30^*th*^ June 2020. We then used the altmetric API to determine the number of media shares as of the 8^*th*^ July 2020. For comparison purposes, we performed the same analysis on non-COVID preprints submitted to arXiv during the same time window. Finally, we performed the same analysis on retracted COVID-19 articles or preprints. The methodology is described in an appendix to this paper, and all the scripts for data extraction and analysis, along with collected data, are available on the OSF repository of the project.

As can be seen in Figure 3, arXiv preprints related to COVID-19 (*n* = 1, 462) were shared more often than preprints on other topics (*n* = 80, 786) submitted during the same period. A similar pattern has also been reported by Fraser et al. [105]. The difference was more pronounced for mentions in the news: whereas 1,066 (1.3%) of non-COVID-19 preprints were mentioned in the news, 156 (10.7%) of COVID-19 preprints were. The numbers for Twitter appear to be much larger than for any other source, probably because of bots that automatically tweet about new preprints, making the comparison on Twitter less relevant. However, the total number of citations (all sources combined) was larger for COVID-19 preprints, with a median of 6 ([*Q*1 − *Q*3] = [2 − 15]) mentions versus 2 ([1 − 5]) for preprints on other research topics. The number of total citations in the media was even higher for preprints available on platforms dedicated to biomedical research. Out of 1,208 COVID-19 preprints found on bioRxiv, the median number of citations was 30 ([15 − 85]) and 444 (36.8%) of these preprints were mentioned at least once in the news. Similarly, out of 4,629 COVID-19 preprints submitted to medRxiv, the median number of citations was 12 ([6 − 36]) and 1,124 (24.3%) of these preprints were mentioned at least once in the news. These findings highlight the increasing trend in preprint sharing during the pandemic, raising concerns about the spread of potentially misleading and unverified data. However, Fraser et al. [105] found that COVID-19 preprints were more commented (on preprint platforms) than non COVID-19 preprints. While this suggests a higher scrutiny for COVID-19 papers, it also illustrate how preprints may encourage scientific debate.

**Figure 3:**
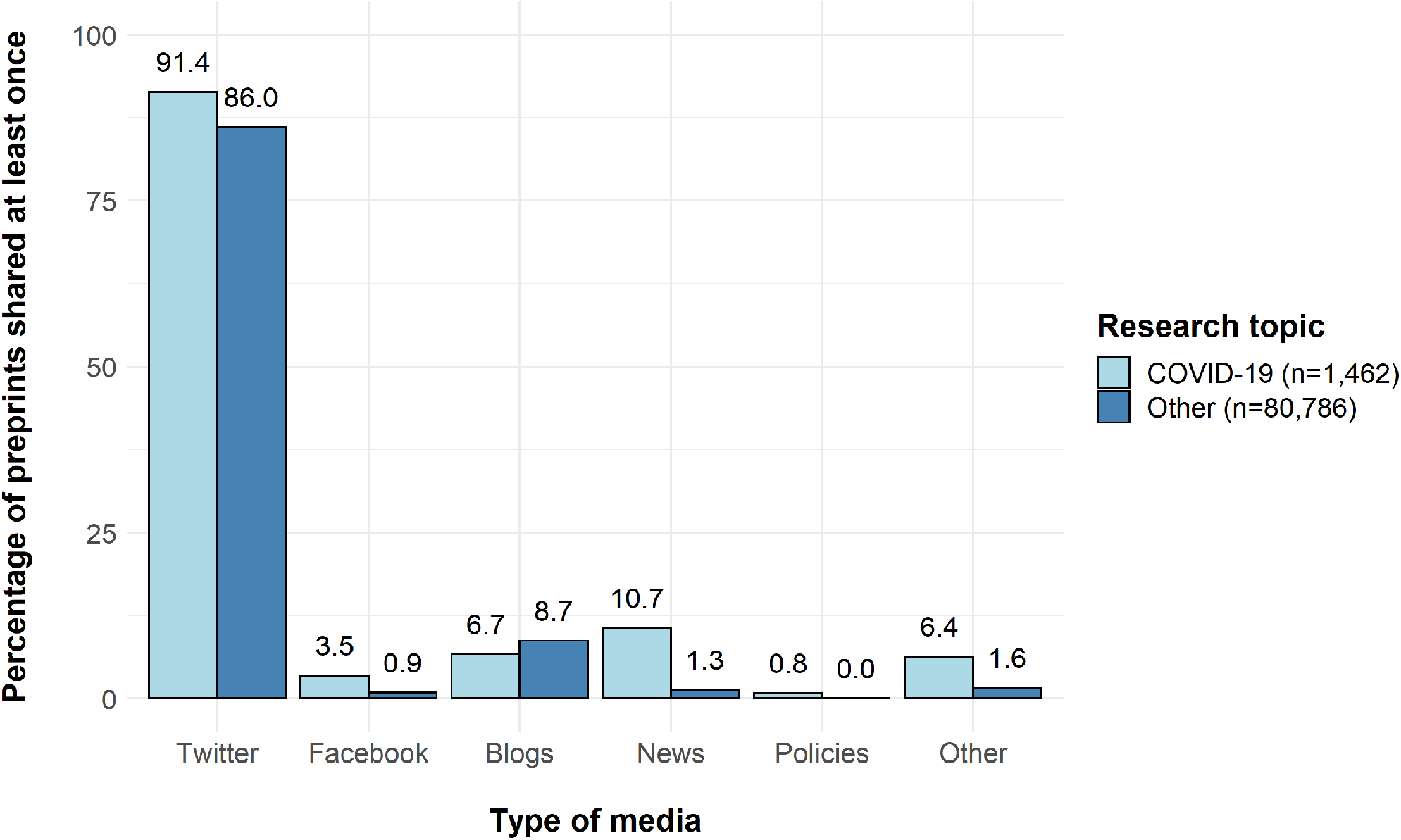
Proportion of arXiv preprints shared in the media, broken down by research topic.

Openly sharing results that have not yet been peer-reviewed can be very damaging if the media and the public take these findings at face value. An outstanding example is the current debate on the effectiveness of hydroxychloroquine as a treatment for COVID-19, which started after the publication of a methodologically flawed preprint [106]. This study quickly caught the attention of the public (1,458 shares in the media including 54 in the news as of 13^*th*^ July 2020), creating a high-demand for the treatment in the absence of valid scientific evidence [107]. This sudden interest contributed to slowing down research conducted on other promising therapeutic strategies. Moreover, the misuse of preprints may discourage researchers from sharing their own in the future, particularly in research areas in which the use of preprints is a quite recent phenomenon, as in medical research (medRxiv was launched in June 2019). Finally, our analysis of retracted papers showed that, among the 6 retracted preprints, the median total of shares was 723 [49 − 2488], emphasizing further the misuse of unverified and, ultimately, invalid findings.

Nevertheless, we would like to emphasize that while peer-review has a key role in maintaining quality, transparency and reproducibility of published articles, it is not sufficient to avoid the publication of flawed studies, and their use by the media and health authorities. Our analysis of retracted papers highlighted the role of some papers, later retracted, in informing public health policy: before the first expression of concern, the study by Mehra et al.[25] was mentioned in 4 policy documents, from 2 sources, including WHO. Besides, this study was the object of 1005 news articles before its retraction, among which 912 were not about the methodological concerns. Similarly, a study investigating the effectiveness of face masks [108] was used in 2 policy documents and was discussed in 132 news outlets.

### 3.2 A call for more reasonable communication

Many science journalists and news editors rely extensively on press releases from institutions [109]. Academic press releases have already been in the spotlight for their impact on the dissemination of exaggerated findings in the news (e.g., [110, 111, 112]) and can therefore directly exacerbate the spread of non-peer-reviewed findings if their communications are based on preprints. Concerns have already been raised about the role of preprints in the communication of science to the public and its potential dangers [113]. Nonetheless, the advantages of preprints for scientific communities are too important to completely give up on preprints [98] and preprints platforms have already implemented warning messages on manuscripts to explain that they have not been peer-reviewed.

An obvious solution to the issue would be to recommend that press releases from research institutions should be made only with respect to peer-reviewed studies, and should be written in collaboration with independent scientists. However, in some cases, it is necessary to report the findings of studies that have not yet been peer-reviewed, if they are expected to benefit the public at large. Should they do so, journalists and news editors must then be encouraged to search for potential conflicts of interest and check the availability of registered information and external peer-reviews to ensure the quality and trustworthiness of their article. Findings from preprints should be communicated with particular caution. Despite the advantages of making headlines simple or even exaggerated [114], journalists and news editors should make sure to accurately convey the inherent degree of uncertainty in scientific studies, which is often quantified and explicitly stated in the original articles. However, such measures are clearly not enough: with the increased productivity pressure on science journalists the fact-checking process needs to be sometimes less thorough [109]. While it seems sometimes difficult [109], journalists and scientists should work together, in particular for scientific results pertaining to public health or that could imply a change of behaviour of the public.

The misuse of preprints by some journalists emphasises the need for high quality journalism training. This issue is not new and has already been pointed out [115, 116, 117], for example during the Ebola crisis [118]. The misuses of research outputs are not limited to preprints but also includes poor quality studies in peer-reviewed and predatory journals. It is clear that a high quality dissemination of scientific information is essential to an appropriate public health response to a crisis such as COVID-19 [119]. The scientific community has addressed this issue by publishing some guidelines for a better dissemination of scientific news to the public [116] and also by fostering bridges between the scientific community and science journalists through exchanges and training [115]. As an example, the French association of science journalists, the <u>Association des journalistes scientifiques de la presse d’information</u>, has launched and funded an exchange program between researchers and journalists. In the UK, the Science Media Centre provides support to news reporters to help them to accurately interpret new findings from publications or press releases [120], by ensuring that journalists, scientists and statisticians work together. Training in science journalism in times of crisis is even more essential and has previously been attempted during the Ebola crisis [118]. During the COVID-19 crisis, the United Nations Educational, Scientific and Cultural Organisation (UNESCO) has organized a series of ‘webinars’ tackling the importance of journalists’ scientific literacy [121]. However, this process is still ongoing and – since many examples of problematic dissemination of scientific news are still seen – some researchers have proposed indicators to evaluate the scientific accuracy of news articles [122]. Some scientists are also grasping these issues by writings blogs directly addressing scientific integrity and exposing examples of bad science in the scientific literature (see e.g., the work of Elisabeth Bik [123]), directly motivating a more responsible science communication.

## Discussion

In addition to previous concerns and investigations of the disruption that the pandemic has caused for research [2], we have found strong evidence of how COVID-19 has impacted science and scientists on several levels. All our findings and the solutions we argued for are summarized in Figure 4. Firstly, we have highlighted the striking scientific waste due to issues in study designs or data analysis. Secondly, we have found that the fast-tracking of peer-reviews on COVID-19 manuscripts, which was needed to give vital treatment directives to health authorities as quickly as possible, led to potentially suspicious peer-reviewing times often combined with editorial conflicts of interest and a lack of transparency of the reviewing process. Thirdly, we highlighted that the lack of raw-data sharing or third-party reviewing has led to the retraction of four major papers and had a direct impact on the study design and conduct of international trials. Finally, we have found evidence of the misuse of preprints in news reports which seem to refer to non-peer-reviewed manuscripts as reliable sources. While we have focused on medical research in the present paper, these issues have also been identified in other research areas, where Open Science principles should also be implemented.

**Figure 4:**
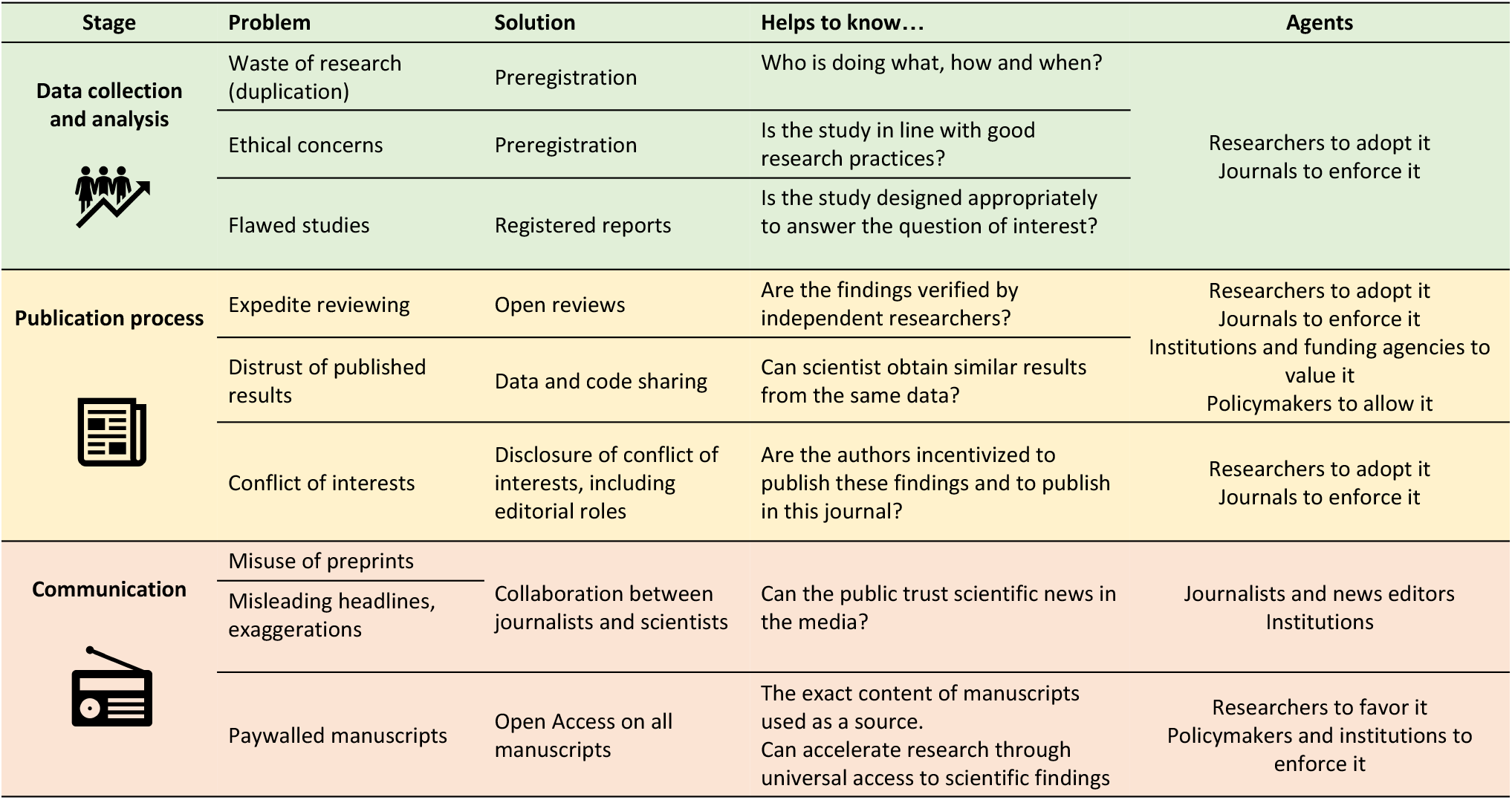
A summary of our findings and proposed solutions.

The Open Science movement promotes more transparency and fairness in the access to scientific communication, the production of scientific knowledge and its communication and evaluation. Looking at the number of publishers removing their paywalls on COVID-19 related research, one might argue that the COVID-19 pandemic has been a catalyst in the adoption of Open Science principles. However, the aforementioned issues paint a more complicated story. The urgency of the situation has led to a partial Open Access policy but with a very opaque peer-review process coupled with a misuse of preprints and raw-data-sharing policies not being enforced. Full adoption of Open Science principles could, however, have saved precious research resources: open peer review would have helped in the detection of the editorial conflicts of interest and made it apparent whether manuscripts were thoroughly reviewed; adoption of registered reports would have strengthened study designs and data analysis plans; proper and monitored use of preprints would have helped the communication of early results between researchers; strengthening of the policies of raw-data sharing or reviewing could have prevented the Surgisphere scandal; and full Open Access might have accelerated the search for solutions to the pandemic both in medical and socio-economic contexts. In addition to this, statistics reviews could have helped to make studies and their results more robust and limit the impact of exaggeration or misinterpretation of results.

It remains, however, that these principles are not enough. The pandemic has highlighted other issues that Open Science cannot solve. For instance, the misuse of preprints by journalists probably stems from the fact that many journalists may not be trained to understand and navigate the complex academic publication system, and some journalist may be seeking sensationalist news headlines. The pandemic has also highlighted the already-existing science-literacy issue [124, 125]. Finally, we cannot exclude that some of the misuses and abuses that we have highlighted are a direct result of the current metric-centered evaluation of research and researchers which has already been shown to lead to questionable research practices in the past and has been the subject of criticism from scientists for decades [42, 126, 127]. Researchers have argued that the adoption of transparency should be coupled with the adoption of a more diverse set of metrics to evaluate researchers [128, 129] or a rejection of metrics altogether [130, 131] to truly limit questionable research practices. A wider adoption of these Open Science Principles cannot be achieved without the endorsement and support of institutions, publishers and funding bodies. International initiatives, such as the Declaration on Research Assessment (DORA), have been put in place to reform the process of research assessment and funding [132], promoting research quality over quantity of outputs. Senior academics have also been identified as key agents in the support of Open Research [133]. For Open Science principles to be clearly and widely adopted, all actors in the scientific community have a role to play: established researchers should encourage a transition to transparent research; institutions and funding agencies should diversify research evaluations; journals, editorial boards, and funding agencies should make all Open Science practices the *de facto* standard for submissions (especially Open Data and registered reports); publishers should strive to make all papers Open Access; and policy-makers and international review boards should consider opening sensible data to reviewers or trusted parties for external validation.

We, as scientific researchers attached to transparency and fairness in the production, communication, use and evaluation of scientific knowledge, hope that this manuscript successfully argues and promotes a faster adoption of all Open Science principles. The call for the changes we argue in this manuscript has been co-signed by 371 researchers (341 of them, or 91.9%, holding a PhD or MD) in the allocated time given by our preregistration: 18 (4.9%) agreed only with the issues we highlighted, 4 (1.1%) agreed only with the solutions we recommended, and the vast majority, 349 (94.1%) agreed with both problems and solutions. We thus hope that this work, supported by so many researchers, will hasten the adoption of transparency as a default in scientific publishing..

## Acknowledgments

We are grateful to Marlon Sidore for fruitful discussions. We also want to thank Matthew Cooper for his helpful feedback on the manuscript.

## Contributions

L.B. and C.L. conceived the study.

L.B., C.L, M.D., H.J., and C.S. extracted the data and performed the analyses.

L.B., N.P.-S., C.Se., P.M., C.Sm., M.D, and C.L. wrote the manuscript.

## Appendix

## Factiva analysis

The Factiva analysis, to find occurences of preprints in the news, was done between January 1^*st*^ 2020 and July 13^*th*^ 2020 with the following query:

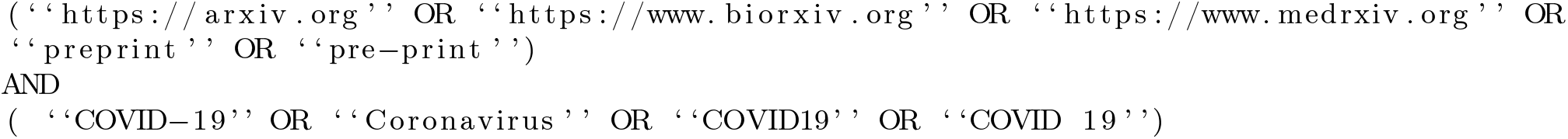

## Altmetric analysis

To further analyse if COVID-19 preprints were used in the news more than preprints are regularly used we conducted an altmetric analysis of all COVID-19 preprints found on arxiv.org, medrxiv.org, and biorxiv.org. We first downloaded all the COVID19-related preprints from these three platforms from the 1st of January 2020 to the 30^*th*^ of June 2020, as well as all preprints from arxiv.org in the same period (to serve as a control group). Duplicates were removed. We then queried the altmetric API for each of these preprints using a Python script to process all entries, find their DOI and query Altmetric with the following command:

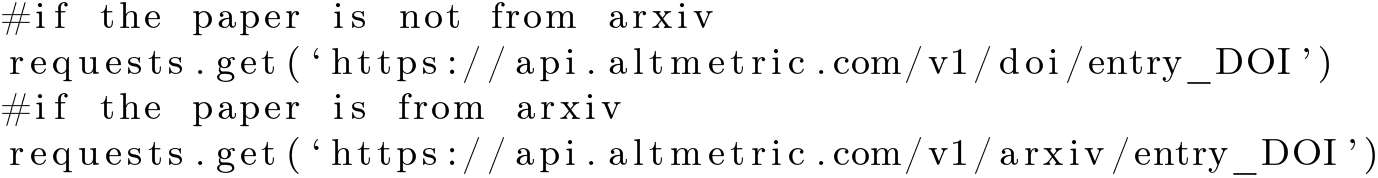

Analysis codes are available on the GitHub repository of the project: https://github.com/lonnibesancon/OpenSciencePandemic

## PubMed Central analysis

To extract the reviewing times, the metadata of 12,682 COVID-19 articles were downloaded on July 7, 2020 from PubMed Central using the query:
“COVID-19”[abstract] OR “COVID-2019“[abstract] OR “severe acute respiratory syndrome coronavirus 2”[Supplementary Concept] OR “severe acute respiratory syndrome coronavirus 2“[abstract] OR “2019-nCoV”[abstract] OR “SARS-CoV-2“[abstract] OR “2019nCoV”[abstract] OR ((“Wuhan”[abstract] AND (“coronavirus”[MeSH Terms] OR “coronavirus”[abstract])) AND (2019/12[PDAT] OR 2020[PDAT])) The reviewing times were extracted from the data using a MATLAB script, available on the OSF repository.

## References

[1] Digital science. (2018-) dimensions [software]. https://app.dimensions.ai (2018, accessed July 20, 2020).

[2] Glasziou, P. P., Sanders, S. & Hoffmann, T. Waste in covid-19 research. BMJ 369, DOI: 10.1136/bmj.m1847 (2020). https://www.bmj.com/content/369/bmj.m1847.full.pdf.

[3] Retracted coronavirus (covid-19) papers. https://retractionwatch.com/retracted-coronavirus-covid-19-papers/ (2020).

[4] Yeo-Teh, N. S. L. & Tang, B. L. An alarming retraction rate for scientific publications on coronavirus disease 2019 (covid-19). Accountability in Research 0, 1–7, DOI: 10.1080/08989621.2020.1782203 (2020). PMID: 32573274, https://doi.org/10.1080/08989621.2020.1782203.

[5] Chalmers, I. & Glasziou, P. Avoidable waste in the production and reporting of research evidence. The Lancet 374, 86–89, DOI: 10.1016/S0140-6736(09)60329-9 (2009).

[6] Allen, C. & Mehler, D. M. A. Open science challenges, benefits and tips in early career and beyond. PLOS Biology 17, 1–14, DOI: 10.1371/journal.pbio.3000246 (2019).

[7] Cockburn, A., Dragicevic, P., Besançon, L. & Gutwin, C. Threats of a replication crisis in empirical computer science. Commun. ACM 63, 70–79, DOI: 10.1145/3360311 (2020).

[8] Munafò, M. R. et al. A manifesto for reproducible science. Nature human behaviour 1, 1–9, DOI: https://doi.org/10.1038/s41562-016-0021 (2017).

[9] Nosek, B. A., Ebersole, C. R., DeHaven, A. C. & Mellor, D. T. The preregistration revolution. Proceedings of the National Academy of Sciences 115, 2600–2606, DOI: 10.1073/pnas.1708274114 (2018). https://www.pnas.org/content/115/11/2600.full.pdf.

[10] Smaldino, P. E., Turner, M. A. & Contreras Kallens, P. A. Open science and modified funding lotteries can impede the natural selection of bad science. Royal Society Open Science 6, 190194, DOI: 10.1098/rsos.190194 (2019).

[11] Masuzzo, P. & Martens, L. Do you speak open science? resources and tips to learn the language. PeerJ Preprints 5, e2689v1, DOI: 10.7287/peerj.preprints.2689v1 (2017).

[12] Watson, M. When will ‘open science’ become simply ‘science’ ? Genome biology 16, 1–3, DOI: 10.1186/s13059-015-0669-2 (2015).

[13] Sarabipour, S. et al. On the value of preprints: An early career researcher perspective. PLOS Biology 17, 1–12, DOI: 10.1371/journal.pbio.3000151 (2019).

[14] Goecks, J., Nekrutenko, A., Taylor, J., Team, G. et al. Galaxy: a comprehensive approach for supporting accessible, reproducible, and transparent computational research in the life sciences. Genome biology 11, R86 (2010).

[15] Miyakawa, T. No raw data, no science: another possible source of the reproducibility crisis. Molecular Brain 13, DOI: https://doi.org/10.1186/s13041-020-0552-2 (2020).

[16] Ross-Hellauer, T. What is open peer review? a systematic review [version 2; peer review: 4 approved]. F1000Research 6, DOI: 10.12688/f1000research.11369.2 (2017).

[17] Besançon, L., Rönnberg, N., Löwgren, J., Tennant, J. P. & Cooper, M. Open up: a survey on open and non-anonymized peer reviewing. Research Integrity and Peer Review 5, 1–11, DOI: https://doi.org/10.1186/s41073-020-00094-z (2020).

[18] Snell, L. & Spencer, J. Reviewers’ perceptions of the peer review process for a medical education journal. Medical Education 39, 90–97, DOI: 10.1111/j.1365-2929.2004.02026.x (2005). https://onlinelibrary.wiley.com/doi/pdf/10.1111/j.1365-2929.2004.02026.x.

[19] Walsh, E., Rooney, M., Appleby, L. & Wilkinson, G. Open peer review: A randomised controlled trial. British Journal of Psychiatry 176, 47–51, DOI: 10.1192/bjp.176.1.47 (2000).

[20] Covid-19 resources for librarians, campuses and health professionals. https://www.elsevier.com/connect/coronavirus-initiatives.

[21] Access covid-19 content across journals, books, and more. https://www.springernature.com/gp/researchers/campaigns/coronavirus#c17669228.

[22] Fraser, N. et al. Preprinting a pandemic: the role of preprints in the covid-19 pandemic. bioRxiv DOI: 10.1101/2020.05.22.111294 (2020). https://www.biorxiv.org/content/early/2020/05/23/2020.05.22.111294.full.pdf.

[23] Pubpeer. https://pubpeer.com/ (2020).

[24] Opensafely. https://opensafely.org/ (2020, accessed July 20, 2020).

[25] Mehra, M. R., Desai, S. S., Ruschitzka, F. & Patel, A. N. Hydroxychloroquine or chloroquine with or without a macrolide for treatment of covid-19: a multinational registry analysis. The Lancet DOI: https://doi.org/10.1016/S0140-6736(20)31180-6 (2020).

[26] M, W. & L, P. Methodological challenges of analysing covid-19 data during the pandemic. BMC Med Res Methodol 20, DOI: 10.1186/s12874-020-00972-6 (2020).

[27] Bik, E. Thoughts on the gautret et al. paper about hydroxychloroquine and azithromycin treatment of covid-19 infections. https://scienceintegritydigest.com/2020/03/24/thoughts-on-the-gautret-et-al-paper-about-hydroxychloroquine-and-azithromycin-treatment-of-covid-19-infections/ (2020).

[28] Rosendaal, F. R. Review of: “hydroxychloroquine and azithromycin as a treatment of covid-19: results of an open-label non-randomized clinical trial gautret et al 2010, doi:10.1016/j.ijantimicag.2020.105949. International Journal of Antimicrobial Agents 106063, DOI: https://doi.org/10.1016/j.ijantimicag.2020.106063 (2020).

[29] Is france’s president fueling the hype over an unproven coronavirus treatment? https://www.sciencemag.org (2020, accessed July 20, 2020).

[30] DeVito N, A. J., Liu M. Covid-19 clinical trials report card: Chloroquine and hydroxychloroquine. https://www.cebm.net/covid-19/covid-19-clinical-trials-report-card-chloroquine-and-hydroxychloroquine/ (2020).

[31] Geleris, J. et al. Observational study of hydroxychloroquine in hospitalized patients with covid-19. N Engl J Med 382, 2411–2418, DOI: 10.1056/NEJMoa2012410 (2020).

[32] H, L. Coronavirus shuts down trials of drugs for multiple other diseases. Nature news (2020).

[33] Gautret, P. et al. Hydroxychloroquine and azithromycin as a treatment of covid-19: results of an open-label non-randomized clinical trial. International Journal of Antimicrobial Agents 105949, DOI: https://doi.org/10.1016/j.ijantimicag.2020.105949 (2020).

[34] Xafis, V., Schaefer, G. O., Labude, M. K., Zhu, Y. & Hsu, L. Y. The perfect moral storm: Diverse ethical considerations in the covid-19 pandemic. Asian Bioethics Review 1 (2020).

[35] Bramstedt, K. A. The carnage of substandard research during the covid-19 pandemic: a call for quality. Journal of Medical Ethics DOI: 10.1136/medethics-2020-106494 (2020). https://jme.bmj.com/content/early/2020/09/09/medethics-2020-106494.full.pdf.

[36] Why should i register and submit results? https://clinicaltrials.gov/ct2/manage-recs/background.

[37] Cockburn, A., Gutwin, C. & Dix, A. Hark no more: On the preregistration of chi experiments. In Proceedings of the 2018 CHI Conference on Human Factors in Computing Systems, CHI ’18, 1–12, DOI: 10.1145/3173574.3173715 (Association for Computing Machinery, New York, NY, USA, 2018).

[38] Reveiz, L. et al. Influence of trial registration on reporting quality of randomized trials: Study from highest ranked journals. Journal of Clinical Epidemiology 63, 1216 – 1222, DOI: https://doi.org/10.1016/j.jclinepi.2010.01.013 (2010).

[39] Dodd, L. E. et al. Endpoints for randomized controlled clinical trials for covid-19 treatments. Clinical Trials 1–11, DOI: 10.1177/1740774520939938 (2020).

[40] Wagenmakers, E.-J., Dutilh, G. & Sarafoglou, A. The creativity-verification cycle in psychological science: New methods to combat old idols. Perspectives on Psychological Science 13, 418–427, DOI: 10.1177/1745691618771357 (2018). PMID: 29961413, https://doi.org/10.1177/1745691618771357.

[41] Wiseman, R., Watt, C. & Kornbrot, D. Registered reports: an early example and analysis. PeerJ 7, e6232, DOI: 10.7717/peerj.6232 (2019).

[42] Chambers, C. D., Feredoes, E., Muthukumaraswamy, S. D. & Etchells, P. Instead of “playing the game” it is time to change the rules: Registered reports at aims neuroscience and beyond. AIMS Neuroscience 1, 4–17, DOI: 10.3934/Neuroscience2014.1.4 (2014).

[43] Bandrowski, A. et al. The resource identification initiative: A cultural shift in publishing. Journal of Comparative Neurology 524, 8–22, DOI: 10.1002/cne.23913 (2016). https://onlinelibrary.wiley.com/doi/pdf/10.1002/cne.23913.

[44] Schulz, K. F., Altman, D. G. & Moher, D. Consort 2010 statement: updated guidelines for reporting parallel group randomised trials. BMJ 340, DOI: 10.1136/bmj.c332 (2010). https://www.bmj.com/content.

[45] Elm, E. v. et al. Strengthening the reporting of observational studies in epidemiology (strobe) statement: guidelines for reporting observational studies. BMJ 335, 806–808, DOI: 10.1136/bmj.39335.541782.AD (2007). https://www.bmj.com/content/335/7624/806.full.pdf.

[46] Society, T. R. Urgent call for registered reports on coronavirus. https://royalsociety.org/blog/2020/03/urgent-call-for-registered-reports-on-coronavirus/ (2020).

[47] LeBlanc, A. G., Barnes, J. D., Saunders, T. J., Tremblay, M. S. & Chaput, J.-P. Scientific sinkhole: The pernicious price of formatting. PLOS ONE 14, 1–7, DOI: 10.1371/journal.pone.0223116 (2019).

[48] Pain, E. Meet octopus, a new vision for scientific publishing. https://www.sciencemag.org/careers/2018/11/meet-octopus-new-vision-scientific-publishing.

[49] Palayew, A. et al. Pandemic publishing poses a new covid-19 challenge. Nature Human Behaviour 1–4, DOI: https://doi.org/10.1038/s41562-020-0911-0 (2020).

[50] Pederson, T. Publishing coronavirology: Peering into peer(less?) review. The FASEB Journal 34, 9825–9827, DOI: 10.1096/fj.202001592 (2020). https://faseb.onlinelibrary.wiley.com/doi/pdf/10.1096/fj.202001592.

[51] Journal of critical care, covid-19 update. https://www.journals.elsevier.com/journal-of-critical-care/covid-19.

[52] Covid-19 articles accepted for fast-track publication in psychological science: Psychological science. https://journals.sagepub.com/page/pss/covid-19.

[53] Full fee waivers and fast-track peer review for covid-19-related manuscripts across all peerj journals. https://peerj.com/blog/post/115284882180/covid-19-full-fee-waivers-fast-track-peer-review/.

[54] The mit press and uc berkeley launch rapid reviews: Covid-19. https://rapidreviewscovid19.mitpress.mit.edu/pub/press-release (2020).

[55] Homolak, J., Kodvanj, I. & Virag, D. Preliminary analysis of covid-19 academic information patterns: a call for open science in the times of closed borders. Scientometrics 124, 2687–2701, DOI: 10.1007/s11192-020-03587-2 (2020).

[56] Aviv, A. Telomeres and covid-19. The FASEB Journal 34, 7247–7252, DOI: 10.1096/fj.202001025 (2020). https://faseb.onlinelibrary.wiley.com/doi/pdf/10.1096/fj.202001025.

[57] on Publication Ethics, C. A short guide to ethical editing for new editors, DOI: 10.24318/cope.2019.1.8.

[58] Risch, H. A. Opinion: Early Outpatient Treatment of Symptomatic, High-Risk Covid-19 Patients that Should be Ramped-Up Immediately as Key to the Pandemic Crisis. American Journal of Epidemiology DOI: 10.1093/aje/kwaa093 (2020). Kwaa093, https://academic.oup.com/aje/advance-article-pdf/doi/10.1093/aje/kwaa093/33585862/kwaa093.pdf.

[59] Fox, M. P. et al. Concerns About the Special Article on Hydroxychloroquine and Azithromycin in High Risk Outpatients with COVID-19 by Dr. Harvey Risch. American Journal of Epidemiology DOI: 10.1093/aje/kwaa189 (2020). Kwaa189, https://academic.oup.com/aje/advance-article-pdf/doi/10.1093/aje/kwaa189/33694717/kwaa189.pdf.

[60] Rosendaal, F. R. Review of: “hydroxychloroquine and azithromycin as a treatment of covid-19: results of an open-label non-randomized clinical trial gautret et al 2010, doi:10.1016/j.ijantimicag.2020.105949. International Journal of Antimicrobial Agents 106063, DOI: https://doi.org/10.1016/j.ijantimicag.2020.106063 (2020).

[61] Machiels, J. D. et al. Reply to gautret et al: hydroxychloroquine sulfate and azithromycin for covid-19: what is the evidence and what are the risks? International Journal of Antimicrobial Agents 106056, DOI: https://doi.org/10.1016/j.ijantimicag.2020.106056 (2020).

[62] Voss, A., Coombs, G., Unal, S., Saginur, R. & Hsueh, P.-R. Publishing in face of the covid-19 pandemic. International Journal of Antimicrobial Agents 56, 106081, DOI: https://doi.org/10.1016/j.ijantimicag.2020.106081 (2020).

[63] Baker, M. 1,500 scientists lift the lid on reproducibility. Nature 533, 452–454 (2016).

[64] Mehra, M. R., Desai, S. S., Kuy, S., Henry, T. D. & Patel, A. N. Retraction: Cardiovascular disease, drug therapy, and mortality in covid-19. n engl j med. doi: 10.1056/nejmoa2007621. New England Journal of Medicine 382, 2582–2582, DOI: 10.1056/NEJMc2021225 (2020).

[65] Organization, W. H. “solidarity” clinical trial for covid-19 treatments. https://www.who.int/emergencies/diseases/novel-coronavirus-2019/global-research-on-novel-coronavirus-2019-ncov/solidarity-clinical-trial-for-covid-19-treatments (2020).

[66] Mahase, E. Covid-19: Who halts hydroxychloroquine trial to review links with increased mortality risk. BMJ 369, DOI: 10.1136/bmj.m2126 (2020). https://www.bmj.com/content/369/bmj.m2126.full.pdf.

[67] Hardwicke, T. E. & Goodman, S. N. How often do leading biomedical journals use statistical experts to evaluate statistical methods? the results of a survey. PLOS ONE 15, 1–12, DOI: 10.1371/journal.pone.0239598 (2020).

[68] The bmj editorial board. https://www.bmj.com/about-bmj/editorial-staff (Accessed 2020-07-15).

[69] Horbach, S. P. & W, H. The ability of different peer review procedures to flag problematic publications. Scientometrics 118, 339–373, DOI: 10.1007/s11192-018-2969-2 (2019).

[70] Pardo, R. et al. Diagnosis and locoregional treatment of patients with breast cancer during the covid-19 pandemic. Revista de Senologia y Patologia Mamaria (2020).

[71] Zenodo. https://zenodo.org/ (2020).

[72] Asapbio. https://asapbio.org/ (2020).

[73] Review commons. https://www.reviewcommons.org/about/ (2020).

[74] Outbreakscience prereview. https://outbreaksci.prereview.org/ (2020).

[75] Morris, T., Dahly, D., Hood, K. & Gates, S. Statistical review of Effect of Dexamethasone in Hospitalized Patients with COVID-19 – Preliminary Report, DOI: 10.5281/zenodo.3928540 (2020).

[76] Bik, E. Post-publication-reviews-on-covid-19-papers. https://scienceintegritydigest.com/2020/03/27/post-publication-reviews-on-covid-19-papers/?

[77] Pucker, B., Schilbert, H. & Schumacher, S. F. Integrating molecular biology and bioinformatics education. Preprints 2018 DOI: 10.20944/preprints201811.0183.v1 (2018).

[78] Jansen, Y., Hornbaek, K. & Dragicevic, P. What Did Authors Value in the CHI’16 Reviews They Received? Proceedings of the 2016 CHI Conference Extended Abstracts on Human Factors in Computing Systems, DOI: 10.1145/2851581.2892576 (2016).

[79] Squazzoni, F., Grimaldo, F. & Marušić, A. Publishing: Journals could share peer-review data. Nature 546, 352 (2017).

[80] McNutt, M. Journals unite for reproducibility. Nature 515, DOI: 10.1038/515007a (2014).

[81] Nichols, T. E. et al. Best practices in data analysis and sharing in neuroimaging using mri. Nature neuroscience 20, 299–303 (2017).

[82] Blumzon, C. F. I. & Pănescu, A.-T. Data Storage, 277–297 (Springer International Publishing, Cham, 2020).

[83] Peiffer-Smadja, N. et al. Machine learning for covid-19 needs global collaboration and data-sharing. Nat Mach Intell 2, 293–294, DOI: http://doi.org/10.1038/s42256-020-0181-6 (2020).

[84] Asendorpf, J. B. et al. Recommendations for increasing replicability in psychology. European Journal of Personality 27, 108–119, DOI: 10.1002/per.1919 (2013). https://onlinelibrary.wiley.com/doi/pdf/10.1002/per.1919.

[85] Begley, C. G. & Ioannidis, J. P. Reproducibility in science: improving the standard for basic and preclinical research. Circulation research 116, 116–126, DOI: 10.1161/CIRCRESAHA.114.303819 (2015).

[86] Fanelli, D. How many scientists fabricate and falsify research? a systematic review and meta-analysis of survey data. PLOS ONE 4, 1–11, DOI: 10.1371/journal.pone.0005738 (2009).

[87] Johnson, A. E. et al. Mimic-iii, a freely accessible critical care database. BSci data 3, DOI: 10.1038/sdata.2016.35 (2016).

[88] Easterbrook, S. M. Open code for open science? Nature Geoscience 7, 779–781, DOI: 10.1038/ngeo2283 (2014).

[89] Johnson, A. E., Stone, D. J., Celi, L. A. & Pollard, T. J. The MIMIC Code Repository: enabling reproducibility in critical care research. Journal of the American Medical Informatics Association 25, 32–39, DOI: 10.1093/jamia/ocx084 (2017). _eprint: https://academic.oup.com/jamia/article-pdf/25/1/32/27901238/ocx084.pdf.

[90] Gilroy, S. P. & Kaplan, B. A. Furthering Open Science in Behavior Analysis: An Introduction and Tutorial for Using GitHub in Research. Perspectives on Behavior Science 42, 565–581, DOI: 10.1007/s40614-019-00202-5 (2019).

[91] Perkel, J. Democratic databases: science on GitHub. Nature 538, 127–128 (2016).

[92] Allen, C. & Mehler, D. M. A. Open science challenges, benefits and tips in early career and beyond. PLoS Biol. 17, e3000246 (2019).

[93] Christensen, G. & Miguel, E. Transparency, reproducibility, and the credibility of economics research. Journal of Economic Literature 56, 920–80, DOI: 10.1257/jel.20171350 (2018).

[94] Wright, D. B. Improving trust in research: Supporting claims with evidence. Open Education Studies 2, 1 – 8, DOI: https://doi.org/10.1515/edu-2020-0106 (2020).

[95] Gymrek, M., McGuire, A. L., Golan, D., Halperin, E. & Erlich, Y. Identifying personal genomes by surname inference. Science 339, 321–324, DOI: 10.1126/science.1229566 (2013).

[96] Wiley’s data sharing policies. https://authorservices.wiley.com/author-resources/Journal-Authors/open-access/data-sharing-citation/data-sharing-policy.html (Accessed 2020-07-15).

[97] Johansson, M. A., Reich, N. G., Meyers, L. A. & Lipsitch, M. Preprints: An underutilized mechanism to accelerate outbreak science. PLOS Medicine 15, 1–5, DOI: 10.1371/journal.pmed.1002549 (2018).

[98] Fry, N. K., Marshall, H. & Mellins-Cohen, T. In praise of preprints. Microbial genomics 5, DOI: 10.1099/mgen.0.000259 (2019).

[99] Kwon, D. How swamped preprint servers are blocking bad coronavirus research. Nature DOI: 10.1038/d41586-020-01394-6 (2020).

[100] Polka, J. K. & Penfold, N. C. Biomedical preprints per month, by source and as a fraction of total literature, DOI: 10.5281/ZENODO.3819276 (2020).

[101] Nih opa isearch covid-19 portfolio. https://icite.od.nih.gov/covid19/search/ (2020).

[102] Johansson, M. A., Reich, N. G., Meyers, L. A. & Lipsitch, M. Preprints: An underutilized mechanism to accelerate outbreak science. PLoS medicine 15, e1002549, DOI: 10.1371/journal.pmed.1002549 (2018).

[103] Huisman, J. & Smits, J. Duration and quality of the peer review process: the author’s perspective. Scientometrics 113, 633–650, DOI: 10.1007/s11192-017-2310-5 (2017).

[104] Abdill, R. J. & Blekhman, R. Meta-research: Tracking the popularity and outcomes of all biorxiv preprints. eLife 8, e45133, DOI: 10.7554/eLife.45133 (2019).

[105] Fraser, N. et al. Preprinting the covid-19 pandemic. bioRxiv DOI: 10.1101/2020.05.22.111294 (2020). https://www.biorxiv.org/content/early/2020/09/18/2020.05.22.111294.full.pdf.

[106] Gautret, P. et al. Hydroxychloroquine and azithromycin as a treatment of covid-19: preliminary results of an open-label non-randomized clinical trial. medRxiv DOI: 10.1101/2020.03.16.20037135 (2020). https://www.medrxiv.org/content/early/2020/03/20/2020.03.16.20037135.full.pdf.

[107] Sattui, S. E. et al. Swinging the pendulum: lessons learned from public discourse concerning hydroxychloroquine and covid-19. Expert Review of Clinical Immunology 0, DOI: 10.1080/1744666X.2020.1792778 (2020). PMID: 32620062.

[108] Bae, S. et al. Effectiveness of surgical and cotton masks in blocking sars–cov-2: A controlled comparison in 4 patients. Annals of Internal Medicine 173, W22–W23, DOI: 10.7326/M20-1342 (2020). PMID: 32251511, https://doi.org/10.7326/M20-1342.

[109] Williams, A. & Clifford, S. Mapping the field: A political economic account of specialist science news journalism in the UK national media. ORCA: Online Research at Cardiff (2008).

[110] Schwartz, L. M., Woloshin, S., Andrews, A. & Stukel, T. A. Influence of medical journal press releases on the quality of associated newspaper coverage: retrospective cohort study. BMJ 344, DOI: 10.1136/bmj.d8164 (2012). https://www.bmj.com/content/344/bmj.d8164.full.pdf.

[111] Sumner, P. et al. The association between exaggeration in health related science news and academic press releases: retrospective observational study. BMJ 349, DOI: 10.1136/bmj.g7015 (2014). https://www.bmj.com/content/349/bmj.g7015.full.pdf.

[112] Sumner, P. et al. Exaggerations and caveats in press releases and health-related science news. PLOS ONE 11, 1–15, DOI: 10.1371/journal.pone.0168217 (2016).

[113] Sheldon, T. Preprints could promote confusion and distortion. Nature 559, 445–446, DOI: 10.1038/d41586-018-05789-4 (2018).

[114] Dumas-Mallet, E., Smith, A., Boraud, T. & Gonon, F. Scientific uncertainty in the press: How newspapers describe initial biomedical findings. Science Communication 40, 124–141, DOI: 10.1177/1075547017752166 (2018).

[115] Meier, K. & Schützeneder, J. Bridging the gaps: Transfer between scholarly research and newsrooms in journalism education—toward an evidence-based practice in an age of post-truth and state of flux. Journalism & Mass Communication Educator 74, 199–211, DOI: 10.1177/1077695819830021 (2019). https://doi.org/10.1177/1077695819830021.

[116] Miranda, G. F., Vercellesi, L. & Bruno, F. Information sources in biomedical science and medical journalism: methodological approaches and assessment. Pharmacological Research 50, 267 – 272, DOI: https://doi.org/10.1016/j.phrs.2003.12.021 (2004).

[117] Peláez, A. L. & Díaz, J. A. Science, technology and democracy: Perspectives about the complex relation between the scientific community, the scientific journalist and public opinion. Social Epistemology 21, 55–68, DOI: 10.1080/02691720601125548 (2007). https://doi.org/10.1080/02691720601125548.

[118] Chalaud, D., Aghan, D., Otindo, V., Bennett, A. & Baldet, T. Ebola: improving science-based communication and local journalism. Lancet 386, 2139 (2015).

[119] La, V.-P. et al. Policy response, social media and science journalism for the sustainability of the public health system amid the covid-19 outbreak: The vietnam lessons. Sustainability 12, DOI: 10.3390/su12072931 (2020).

[120] Science media centre. https://www.sciencemediacentre.org/ (2020, accessed July 20, 2020).

[121] Scientific journalism on covid-19 time. https://en.unesco.org/events/scientific-journalism-covid-19-time (2020).

[122] Smeros, P., Castillo, C. & Aberer, K. Scilens: Evaluating the quality of scientific news articles using social media and scientific literature indicators. In The World Wide Web Conference, WWW ’19, 1747–1758, DOI: 10.1145/3308558.3313657 (Association for Computing Machinery, New York, NY, USA, 2019).

[123] Science integrity digest. https://scienceintegritydigest.com/ (2020).

[124] Kennedy, B. & Hefferon, M. What americans know about science: Science knowledge levels remain strongly tied to education; republicans and democrats are about equally knowledgeable. Pew Research Center (2019).

[125] Vraga, E. K., Tully, M. & Bode, L. Empowering users to respond to misinformation about covid-19. Media and Communication 8, 475–479, DOI: https://doi.org/10.17645/mac.v8i2.3200 (2020).

[126] It is time to fix research evaluation. https://opensciencemooc.eu/evaluation/2019/10/15/solve-research-evaluation/ (2019).

[127] Nosek, B. A., Spies, J. R. & Motyl, M. Scientific utopia: Ii. restructuring incentives and practices to promote truth over publishability. Perspectives on Psychological Science 7, 615–631, DOI: 10.1177/1745691612459058 (2012).

[128] Barrett, L. F. The publication arms race. APS Observer 32(2019).

[129] Jeschke, J. M., Lokatis, S., Bartram, I. & Tockner, K. Knowledge in the dark: scientific challenges and ways forward. FACETS 4, 423–441, DOI: 10.1139/facets-2019-0007 (2019). https://doi.org/10.1139/facets-2019-0007.

[130] Crous, C. J. The darker side of quantitative academic performance metrics. South African Journal of Science 115, 1 – 3, DOI: http://dx.doi.org/10.17159/sajs.2019/5785 (2019).

[131] Lawrence, P. A. The mismeasurement of science. Current Biology 17, R583–R585, DOI: https://doi.org/10.1016/j.cub.2007.06.014 (2007).

[132] for Cell Biology., A. S. Dora. declaration on research assessment. https://sfdora.org/read/ (2012).

[133] Kowalczyk, O., Lautarescu, A., Blok, E., Dall’Aglio, L. & Westwood, S. J. What senior academics can do to support reproducible and open research: a short, three-step guide, DOI: 10.31234/osf.io/jyfr7 (2020).

